# The contribution of electrostatics to hydrogen exchange in the unfolded protein state

**DOI:** 10.1101/2021.02.22.432104

**Authors:** Rupashree Dass, Enrico Corlianò, Frans A. A. Mulder

## Abstract

Although electrostatics have long been recognized to play an important role in hydrogen exchange (HX) with solvent, the quantitative assessment of its magnitude in the unfolded state has hitherto been lacking. This limits the utility of HX as a quantitative method to study protein stability, folding and dynamics. Using the intrinsically disordered human protein α-synuclein as a proxy for the unfolded state, we show that a hybrid mean-field approach can effectively compute the electrostatic potential at all backbone amide positions along the chain. From the electrochemical potential a fourfold reduction in hydroxide concentration near the protein backbone is predicted for the C-terminal domain, a prognosis that is in direct agreement with experimentally-derived protection factors from NMR spectroscopy. Thus, impeded HX for the C-terminal region of α-synuclein is not the result of intramolecular hydrogen bonding and/or structure formation.

## Introduction

Hydrogen exchange substitution was among the earliest characterization methods to demonstrate the globular nature of proteins (1) and has been instrumental in demonstrating the presence of partial protein unfolding, the existence of a hierarchy of protein states between the unfolded and folded states of proteins, and for the determination of the local stability of protein structure to unfolding (2–4). Typically, HX measurements follow the rate at which labile backbone amide hydrogens (protium, ^1^H) exchange with deuterium (^2^H) upon dissolution into D_2_O. Alternatively, HX can be probed by transfer of magnetization saturation from the water in NMR experiments such as CLEANEX-PM (5). Chemically, the rate of exchange depends on various factors, including burial, hydrogen-bond formation, amino acid sequence composition and external variables (such as pressure, temperature, and ionic strength) (6, 7). Protein structuration will act to slow HX, and this deceleration is routinely expressed by ‘protection factors’ (PFs); The PF is defined as the ratio of the ‘intrinsic’ exchange rate, k_intr_, in the fully unprotected, solvent-accessible state to the observed rate, k_obs_. Reduced HX rates may also arise from other mechanisms, including inductive effects from neighboring side chains (8, 9), long-range electrostatic interactions (10) and the local distribution of charged residues (11). For example, amide HX rates are very sensitive to the amino acid sequence (9), and these (mainly inductive) effects are due to the stereo-electronic properties of side chains to the left and right side of the amide group. As ‘intrinsic’, sequence-specific reference exchange rates for the protein sequence under consideration are typically not available, intrinsic amide HX rates are computed by proxy from data tabulated for host peptides, such as poly-D,L-alanine (8). ‘Intrinsic’ exchange rates are thus straightforwardly, but only approximately, predicted from the primary structure by con-sideration of the identity of the side chains bracketing each of the amide hydrogens in the sequence using a lookup table. Protection factors have a simple interpretation: For example, PF = 4 means that the amide proton is protected PF/(1 + PF) = 80% of the time. Therefore, HX is an attractive method to study the local structural stability in proteins, including the presence of structural motifs or domains in disordered regions (12–14).

However, to date, a quantitative description of the influence of electrostatics on HX in the unfolded state has been lacking, and this continues to limit the precision of protection factors and their meaningful interpretation. In this paper we compute the electric potential along the unfolded protein chain from sequence using a mean-field approach. As a proxy for the unfolded state, we used human α-synuclein, an intrinsically disordered protein (IDP) of 140 amino acids, which is one of the most unfolded proteins currently characterized (15). Of interest for this work, the protein harbors three regions with very distinct charge patterning; an N-terminal domain (1-61) rich in positively and negatively charged amino acids and with a small net positive charge, a central region (62-95; also known as the NAC region) that is devoid of charged amino acids with two exceptions, and a highly acidic C-terminal tail (96-140). Using the electrochemical potential, the concentration reduction of catalytic hydroxide concentration near the polyamide backbone was calculated to be fourfold, and shown to match with experimental observation.

## Materials and Methods

### Sample preparation

For hydrogen exchange measurements, 10 mg ^13^C/^15^N-labelled α-synuclein (Giotto Biotech, Sesto Fiorentino, Italy) was dissolved in 550 μl 25 mM Tris buffer pH 9, to which 50 μl D_2_O and 10 μl 50 mM 4,4-dimethyl-4-silapentane-1-sulfonic acid (DSS) were added. For co-solute PRE measurements, 2D ^1^H-^15^N HSQC experiments were acquired. For these measurements three samples were made, each containing 200 μM ^13^C/^15^N-labeled α-synuclein, 25 mM phosphate buffer pH 7.3 and 10% D_2_O. Gadoteric acid (C_16_H_25_GdN_4_O_8_) (sample A), or gadoteridol: (C_17_H_29_GdN_4_O_7_) (sample B) were added to 1 mM complex concentration. Sample C (no added PRE agent) served as a diamagnetic reference.

### Experimental NMR set-up

Experiments were recorded on a Bruker spectrometer at ^1^H frequency of 950 MHz equipped with a cryogenically-cooled triple resonance probe. Chemical shifts were referenced with respect to DSS. Backbone assignments were taken from BMRB (https://bmrb.io/) entry 18857, and verified by conducting a 3D HNCO experiment at 283 K and ^1^H-^15^N HSQC experiments at 283 K, 288 K and 298 K for the sample at pH 7.52. 90° Pulse lengths were 10 μs (^1^H), 10 μs (^13^C) and 33 μs (^15^N).

For determination of hydrogen exchange, a 2D CON experiment was adapted to include a DÉCOR module (16). The pulse sequence is available for download from www.protein-nmr.org. Experiments were performed at 298 K, pH 9. 2D data matrices consisted of 512(^13^C) x 256(^15^N) points (real + imaginary). The ^15^N CPMG block was run with 2, 4, 6, 8, or 12 π pulses in a constant time of 20 ms. The spectral widths were 3,939 Hz (^13^C) and 2,310 Hz (^15^N), respectively. Each experiment was run with 8 scans, taking 5.5 hours per 2D dataset. A reference experiment was run at each pH and temperature, in which WALTZ-65 decoupling with RF power of 5.0 kHz was inserted on the ^1^H channel during the CPMG period.

Gradient sensitivity-enhanced ^1^H-^15^N HSQC experiments were performed for measuring PRE effects at 283 K. The experiments were acquired with decoupling during acquisition and with 2,048 (^1^H) and 256 (^15^N) time domain points. The spectral widths were 15,243 Hz (^1^H) and 3,177 Hz (^15^N). The addition of paramagnetic complex did not cause changes in peak positions, indicating that it did not cause any change in structure or that it bound to the protein. All NMR data were processed by NMRPipe (17) and analyzed by Sparky (18).

### Determination of HX rate constants and protection factors

Peak Intensities of the CON spectra (extracted using Sparky) were used to calculate the rate constant using peaks for which the SNR was larger than 5. The calculations followed published work (19, 20). Protection factors were calculated from the relation PF = k_intr_/k_obs_, where k_intr_ was calculated from the protein sequence using the SPHERE server with default parameters (https://protocol.fccc.edu/research/labs/roder/sphere/sphere.html).

### Determination of amide proton temperature coefficients

Amide proton chemical-shift temperature coefficients were calculated using published data that was recorded at 278, 288, 293, 298, and 303 K (BMRB entry 18857). In Figure 2, a dashed line is drawn at –4.5 ppb/K, which demarcates the approximate border found to correspond to hydrogen bond formation in folded proteins (21, 22).

### Calculation of the electrostatic potential

An equation for the energy based on analytical form (23) was used to calculate the potential as the energy divided by unit charge, using the hybrid mean-field approach of Tamiola *et al*. (24). The potential at each backbone position was evaluated using T=298 K; I= 0.025 M. The effective bond length for the Gaussian Chain model was 7.5 Å, and the shift for the distance to the side chain was 10 Å. At pH 9 the charges were defined as: “D”: −1.0, “E”: −1.0, “H”: 1.0, “K”: 1.0, “R”: 1.0. The termini were charged.

## Results and Discussion

Although experimental HX rates for α-synuclein have been reported in the literature using cross-saturation (i.e. by CLEANEX-PM), we were concerned about possible artifacts stemming from cross-relaxation (25) or exchange from the hydroxyl groups of serine and threonine side chains (26). We therefore developed an alternative approach to measure HX that relies on exchange-induced incoherent dephasing of two-spin correlations (16, 27), which is immune to relayed magnetization transfer artifacts. To be generally applicable to unfolded proteins and IDPs, where backbone amides may exchange so rapidly that they are not detectable at the amide hydrogen (28), we designed a proton-less 2D CON pulse sequence that employs ^13^C detection (29) and relies on the highly favorable dispersion of backbone ^15^N and ^13^C’ chemical shifts in non-folded proteins. The experiment (20) contains a CPMG (30, 31) block on the ^15^N channel, where the density operator proportional to 2C_z_N_y_ is allowed to partially evolve into 4C_z_N_x_H_z_, generating an admixture that depends on the inter-pulse spacing chosen by the experimenter. As chemical exchange will exclusively act to annihilate 4C_z_N_x_H_z_ by spin decorrelation (27, 32), exchange-dependent signal loss will ensue. As the method does not rely on the detection of the amide proton, we were able to detect all fast exchanging residues of α-synuclein that otherwise disappear from HSQC spectra at high pH (26).

A series of experiments (A) was conducted by changing the number of CPMG π pulses but keeping the total CPMG time constant, to produce different admixtures of 2C_z_N_y_ and 4C_z_N_x_H_z_. To obtain a reference signal which does not contain the exchange contribution, a second experiment (B) was conducted with WALTZ-65 decoupling on ^1^H throughout the CPMG block. Exchange rates were determined by fitting computed ratios (A/B) to the experimentally obtained ones (16, 27) using published protocols (20). The exchange rates are shown in Figure 1a. Significant variation is observed along the sequence, with a marked plunge at the C-terminus, in agreement with earlier observations (26, 33, 34). To account for the expected variation in intrinsic amino-acid sequence-dependent exchange rates (9), the measured exchange rates were converted to PFs using the program SPHERE (35, 36) as shown in Figure 1b. This greatly reduces the spread in the data for the region 1–100, where PFs vary around one – meaning a complete lack of order. In marked contrast, the C-terminus displays values much larger than one, which might be taken to mean that residual structure is present in the C-terminal domain (33).

**Figure 1.**
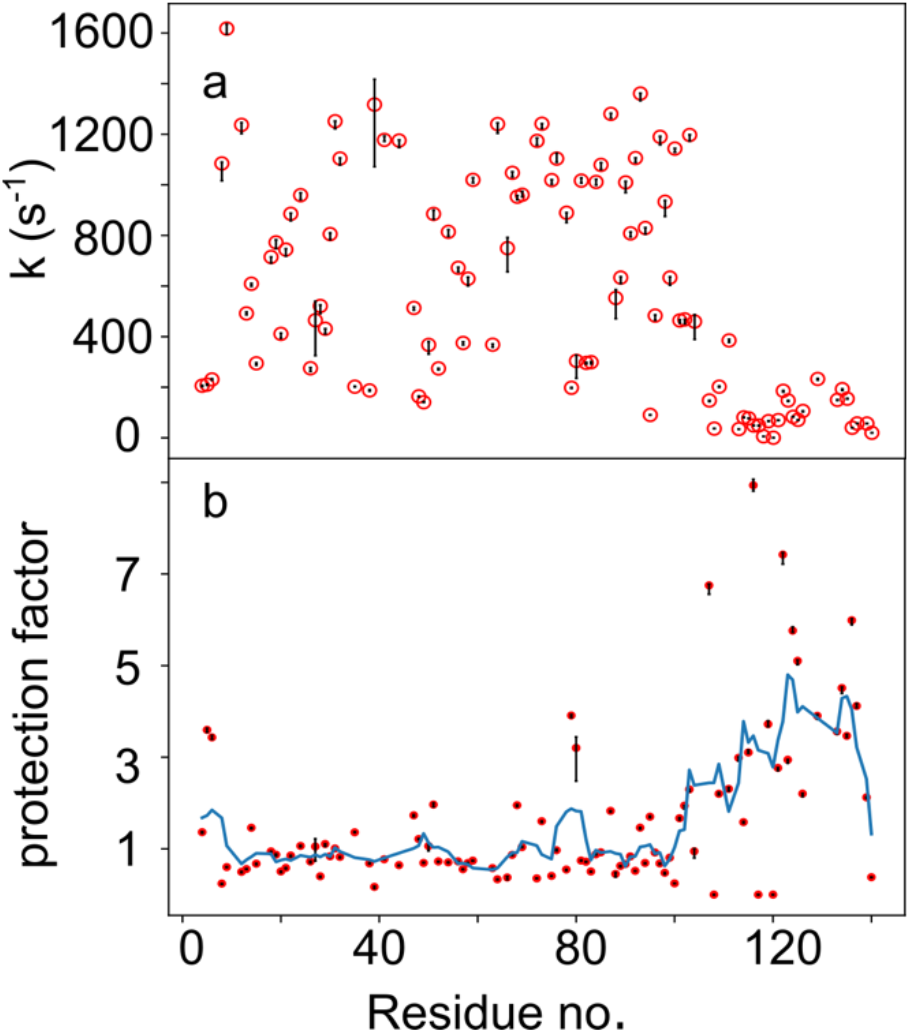
Backbone amide hydrogen exchange of α-synuclein. (a) Exchange rates with solvent at pH 9, 298 K. (b) Hydrogen exchange protection factors. To guide the eye, a moving aver-age of length 5 is shown as solid line in panel (b).

As this insinuation is at variance with analyses of NMR chemical shifts and scalar couplings for the carboxy-terminus, we sought to evaluate the presence of hydrogen bonds by amide hydrogen temperature coefficients, (dδ/dT); In folded proteins, comprehensive analyses have shown that impeded HX due to hydrogen-bond formation is accompanied by reduced temperature coefficients, such that dδ/dT < –4.5 ppb/K provide strong support for H-bonding (21, 22). Amide hydrogen temperature coefficients were determined from deposited chemical shift for α-synuclein at several temperatures (BMRB entry 18857), and are shown in Figure 2. For the most part, values fluctuate around –6 to –7 ppb/K, typical of unstructured regions. Although possibly somewhat reduced values are seen for regions 30-50 and 120-140, these cannot explain the large PFs observed. Consequently, this data agrees with a broad body of evidence that suggests an absence of persistent interactions in the monomeric protein, with only subsidiary weak electrostatic interactions existing between the N– and C–termini.

**Figure 2.**
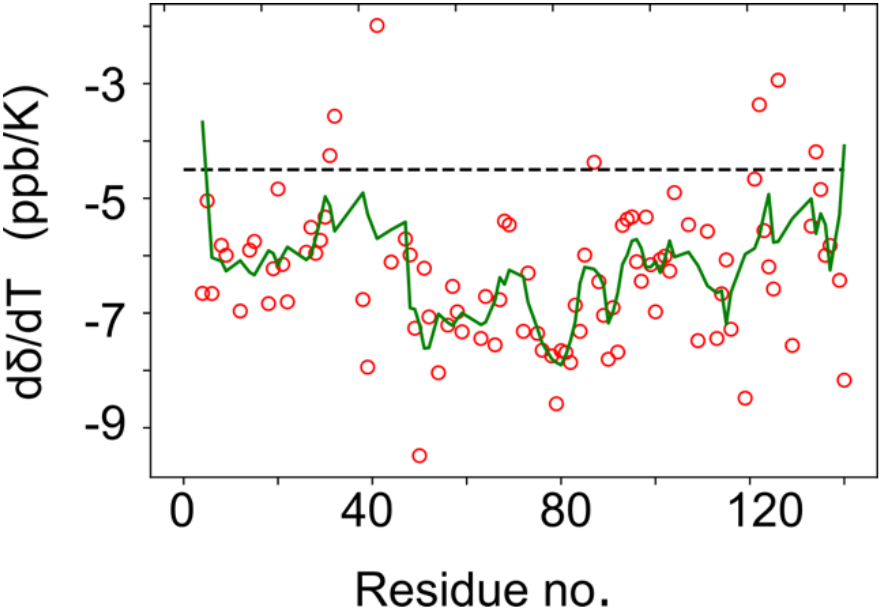
NMR chemical shift temperature coefficients for the amide hydrogens of α-synuclein. To guide the eye, a moving average of length 5 is shown as solid line.

At the same time, the preponderance of Asp and Glu residues at the C-terminus of α-synuclein will create a negative potential around the protein chain. Although the effect of the electrostatic surface potential at the protein-solvent interface on hydrogen exchange has been qualitatively discussed in the past (37, 38), an experimental demonstration of exclusion of charged species near the backbone has hitherto been lacking. To demonstrate its reality, we compared co-solute paramagnetic relaxation enhancement (sPRE) (39, 40) induced by two paramagnetic Gd(III) complexes that are physicochemically equivalent, except for their net charge: Whereas gadoteric acid is negatively charged, gadoteridol is neutral. When added to the protein solution, dipolar interactions with the unpaired lanthanide electrons (S = 7/2) increases the relaxation rates of the protein nuclei with a 1/r^6^ distance dependence, which causes broadening of resonances for nuclei in close proximity. We compared changes in peak broadening for solutions with neutral and negatively charged co-solute, where any region of the protein that preferentially excludes the charged ionic species will display increased intensity. Figure 3a shows the ratio of peak heights when the negative complex is added to the sample, compared to when a neutral complex is added to the sample. Whereas the neutral complex affected backbone amides more evenly, in accordance with published investigations using Ni(II) and Fe(III) complexes (41, 42), the negatively charged complex caused less line broadening at the C-terminus, in accord with electrostatic repulsion

**Figure 3.**
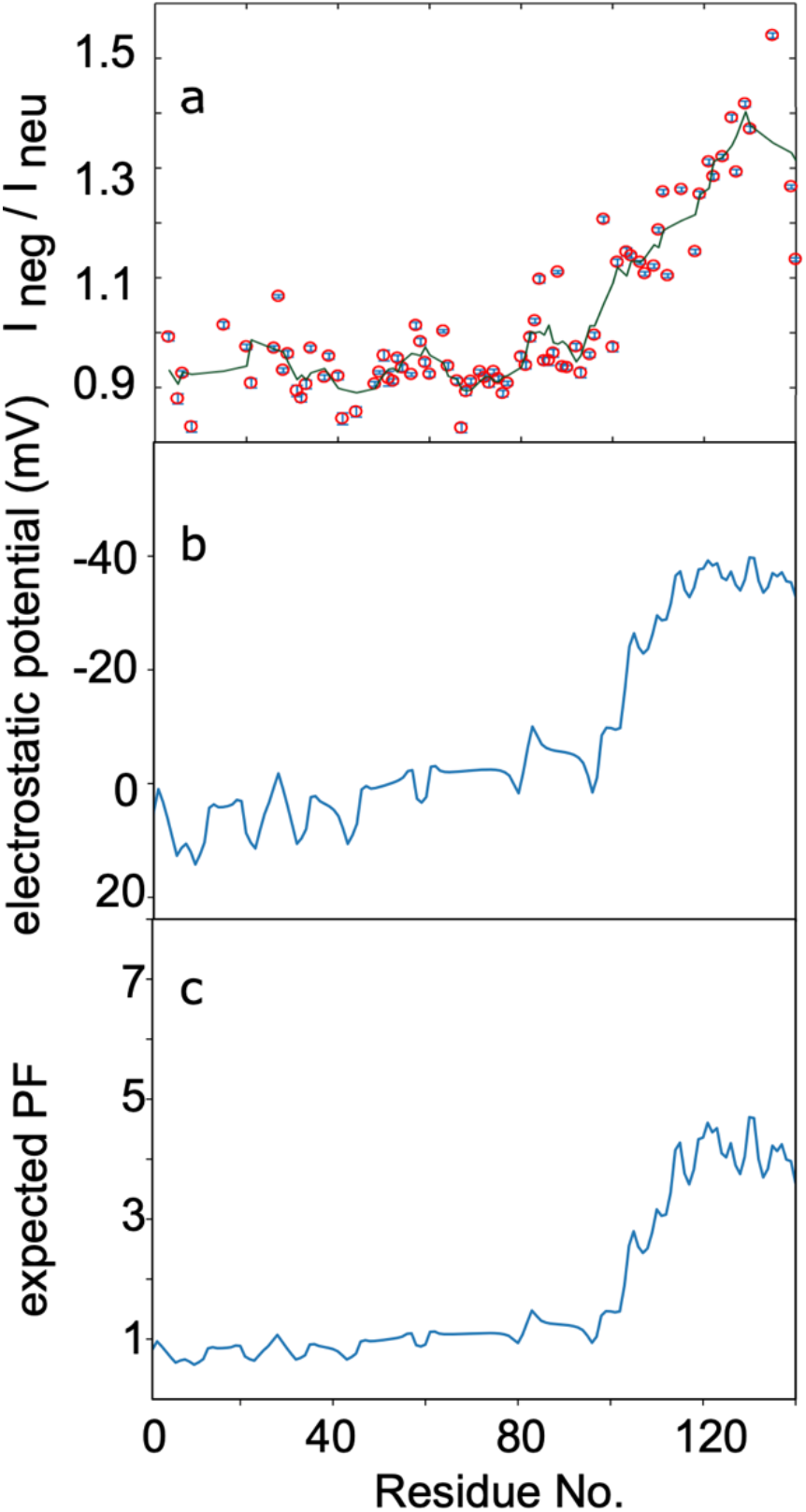
Correspondence between (a) reduced access to negatively charged paramagnetic chelate, (b) calculated electrostatic potential and (c) predicted protection factors for α-synuclein. To guide the eye, a moving average of length 5 is shown as solid line in panel a.

A quantitative understanding of ion depletion can be gleaned by computing the electrostatic potential at the protein backbone. Using the Debije-Hückel approximation for a Gaussian polyelectrolyte protein chain (23, 24), we computed the electrostatic potential at each amino acid in the α-synuclein backbone as:

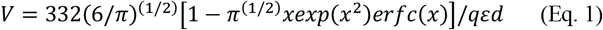

This computed potential is shown in Figure 3b. It matches very well with the observed reduction in HX, as well as with the exclusion of the negatively charged PRE agent. Since hydroxide ions in solution are the species (43) that catalyze backbone amide HX above pH 5, a lower concentration of OH^−^ in the vicinity of the polypeptide chain would result in a concomitant reduction in exchange kinetics, in agreement with the observations made for gadoteric acid.

In thermodynamic terms, to maintain a constant electrochemical potential at the polypeptide backbone at equilibrium, the electrostatic interaction energy acts to lower the local hydroxide concentration (44):

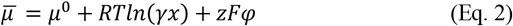

Here μ^0^ is the chemical potential of the pure substance in the standard state (i.e. at reference absolute temperature T and pressure p, where all interactions have been extinguished), R is the universal gas constant, γ is the activity constant, x is the mole fraction of ions (the activity a = γ·x), z is the charge on the ion, F is the Faraday constant and φ is the electric potential. Thus, it is possible from equation (2) to use the computed potential φ to estimate the concentration change of hydroxide at the protein backbone, and therewith predict protection factors. Identifying that the PF is proportional to the fold-change in hydroxide concentration, rearrangement of Eq (2) gives:

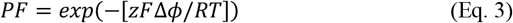

Thus, from the calculated electric potential, a quantitative prediction of the retardation effect on base-catalyzed HX can be made. The computed PFs for α-synuclein are shown in Figure 3c and the trend is in good agreement with the experimental data (Figure 1b). The scatter that is observed in the experimentally derived PFs is wholly a consequence of shortcomings in the modelling of the intrinsic rate constants, and suggests that there is considerable room for improvement in the modelling of intrinsic HX rates for polypeptides. Although the model for calculating the electrostatics is approximate, the agreement attained here is at a level that is similar to predictions of p*K*_a_ constants in unfolded polypeptides (23, 24). The observed correspondence is therefore highly encouraging, and should allow more realistic ensemble-based calculations to be tested in the future.

In an early study, Alexandrescu and co-workers demonstrated (26) that amide hydrogens exchanged rapidly with solvent, confirming the unfolded nature of α-synuclein. They also explained why NMR signals were lost in in-cell ^15^N-^1^H HSQC spectra of α-synuclein upon raising the temperature and pH, providing a first demonstration of the persistence of its disordered state *in vivo* (45, 46). Using CLEANEX-PM HX (5), they noticed that the C-terminal region exchanged slower than the rest of the protein, and sought this explanation in the possibility that the large number of polar side chains (Thr and Ser in particular) would lead to enhanced exchange for the N-terminus relative to the C-terminal region. They also tested whether electrostatics might play a role by the addition of 300 mM NaCl, and found this to (i) enhance exchange rates and (ii) erase the difference in rates between the N- and C-termini. A “change in the effective pH in the vicinity to the protein compared to bulk solution” and “screening the protein from the reactive H_3_O^+^ and OH^−^ ions that catalyze exchange” were suggested as two possible explanations, but neither hypothesis was tested in their work. At a minimum, structural differences of the protein between the two conditions could be excluded as an explanation, as the addition of salt did not induce chemical shift changes. Subsequently Okazaki *et a*l. performed a more extensive study of HX for α-synuclein at 100 mM NaCl by CLEANEX-PM, recording data at various pH values and experimental mixing times (33). They consistently found slower exchange in the C-terminal region at all pH values studied. When converting their data to protection factors, a pattern emerged with values around 1 (indicating full solvent exposure) for the region 30– 100, whereas relatively scattered values were obtained for the amino-terminus. Interestingly, in agreement with the Croke study (26), increased PFs were obtained for the C-terminal domain. Although protection factors are expected to exhibit some scatter due to the limited accuracy of computed intrinsic rates from the lookup tables (26, 47) the pervasive increase observed for the C-terminus was well-reproduced in the two studies. Although Okazaki *et al*. acknowledged that exchange may be hindered by the presence of negative charge, they concluded that interaction between the N- and C-termini is the dominant factor for retardation (33). The existence of long-range interactions has also been suggested by PRE studies (48, 49) using the introduction of spin labels at cysteine residues. Although misincorporation (50) of Cys at position 136 may have potentially compromised these studies by leading to unintentional signal loss in the C-terminus, weak electrostatic interaction between the N- and C-termini are supported by other data (51, 52) and have been suggested to persist in the cellular milieu (45). At the same time, monomeric α-synuclein is intrinsically disordered along its entire sequence at the local level as judged from NMR chemical shifts (15) and scalar couplings (53). As it stands, a cogent explanation that encompasses both the lack of structure formation of α-synuclein as well as impeded hydrogen exchange for the C-terminus is lacking.

The combination of electrostatic modelling of the unfolded state, coupled to computation of the change in electrochemical potential now shows that the paradox is resolved by a quantitative consideration of repulsion of catalytic hydroxide ions by acidic protein side chains. The fact that electrostatics due to local amino acid biases can have such significant effects on intrinsic HX rates has significant implications for the structural and energetic interpretation of protection factors. As observed in this study, even for an IDP that is qualified as 97% disordered (15), intrinsic exchange rates calculated from the Bai/Englander tables display a wide variation, and computed protection factors range from 0.1–10. The concurrent investigation of amide hydrogen chemical shift temperature coefficients shows that even this large variation cannot be used to infer a dependable level of protection due to hydrogen bonding.

As one of the main application areas of PFs is the computation of residue-specific stabilities to unfolding, this means that ∆∆G values in protein stability studies from NMR- or MS-based HX are in error by 1.7 kJ/mol for every factor two that intrinsic rates are over- or underestimated. What is more, for regions of relatively high charge density – and in particular for studies performed at low ionic strength – rates in the unfolded form will show large, systematically skewed values that would lead to sizeable deviations. The lack of better, more appropriate models for (locally) disordered, exchange-competent states that are operational in HX, clearly limits the interpretation accuracy of stability data. By the same token, moderate variations of ∆∆G values observed for structural elements do not necessarily mean that they do not unfold cooperatively.

A comprehensive model for computing intrinsic exchange rates of polypeptide amide backbone hydrogens that can account for electrostatic effects offers exciting prospects to utilize HX measurements as quantitative proxies for determining the electrostatic potential around intrinsically disordered proteins. Such methodologies would also be valuable for studies of protein liquid-liquid phase separation, where electrostatics play a key role in the formation of coacervates (54, 55). Studies along these lines are in progress in our laboratory.

## Conclusion

It is shown here that the concentration of anions in the vicinity of an acidic disordered (unfolded) protein is reduced in a way that accurately mirrors the electrostatic potential. This quantity was efficiently calculated and predicted the fourfold increase in protection from hydrogen exchange that was observed in experiment. These results demonstrate that a local drop in the hydroxide concentration at the protein backbone amide is the single dominant factor to explain the reduced HX rates for the C-terminal region of α-synuclein. In other words, impeded hydrogen exchange at the acidic tail of human α-synuclein is a direct experimental manifestation of the electrochemical potential.

## Acknowledgements

Access to the NMR spectrometers at the Danish Center for Ultrahigh-Field NMR Spectroscopy (Ministry of Higher Education and Science grant AU-2010-612-181) is gratefully acknowledged.

## Notes

### Competing Interest Statement

The authors have declared no competing interest.

