## Supplementary Material for "The contribution of electrostatics to hydrogen exchange in the unfolded protein state"

**Table S1**. Experimental and computed parameters for a-synuclein: Measured hydrogen exchange rate, protection factor, amide hydrogen temperature coefficient, signal intensity ratio in the presence of negative divided by neutral PRE agent, calculated electrostatic potential, and calculated protection factor.

| peptide bond | rate ($s^{-1}$) | measured PF | Temp. coeff. (ppb/K) | $I_{neg}/I_{neu}$ | electrostatic potential (mV) | expected PF |
|---|---|---|---|---|---|---|
| D2N-M1C | - | - | - | - | 1.381 | 0.817 |
| V3N-D2C | - | - | -7.222 | 0.992 | 3.657 | 0.948 |
| F4N-V3C | 206.017 | 1.364 | -6.657 | - | 6.587 | 0.867 |
| M5N-F4C | 210.233 | 3.596 | -5.043 | 0.879 | 9.820 | 0.774 |
| K6N-M5C | 230.763 | 3.432 | -6.659 | 0.926 | 13.137 | 0.682 |
| G7N-K6C | - | - | -5.689 | - | 11.792 | 0.600 |
| L8N-G7C | 1084.645 | 0.242 | -5.819 | 0.828 | 11.057 | 0.632 |
| S9N-L8C | 1617.746 | 0.602 | -5.992 | - | 12.485 | 0.650 |
| K10N-S9C | - | - | -6.968 | - | 14.719 | 0.615 |
| A11N-K10C | - | - | -7.395 | - | 13.032 | 0.564 |
| K12N-A11C | 1237.161 | 0.497 | -6.965 | - | 10.852 | 0.602 |
| E13N-K12C | 491.806 | 0.559 | - | - | 4.815 | 0.655 |
| G14N-E13C | 609.032 | 1.460 | -5.903 | - | 4.233 | 0.829 |
| V15N-G14C | 294.599 | 0.675 | -5.751 | 1.014 | 4.733 | 0.848 |
| V16N-V15C | - | | -8.024 | - | 4.726 | 0.832 |
| A17N-V16C | - | - | -7.000 | - | 4.590 | 0.832 |
| A18N-A17C | 714.763 | 0.943 | -6.835 | - | 4.319 | 0.836 |
| A19N-A18C | 772.548 | 0.872 | -6.227 | - | 3.521 | 0.845 |
| E20N-A19C | 410.93 | 0.506 | -4.838 | 0.974 | 3.768 | 0.872 |
| K21N-E20C | 744.21 | 0.585 | -6.151 | 0.908 | 9.413 | 0.864 |
| T22N-K21C | 885.606 | 0.854 | -6.805 | - | 11.106 | 0.693 |
| K23N-T22C | - | | -6.724 | - | 12.153 | 0.649 |
| Q24N-K23C | 959.216 | 1.063 | - | - | 9.009 | 0.623 |
| G25N-Q24C | - | - | -4.395 | - | 6.140 | 0.704 |

| | | | | | | |
|---|---|---|---|---|---|---|
| V26N-G25C | 275.064 | 0.723 | -5.941 | 0.972 | 4.087 | 0.787 |
| A27N-V26C | 464.756 | 1.050 | -5.505 | 1.066 | 1.535 | 0.853 |
| E28N-A27C | 521.521 | 0.399 | -5.959 | 0.932 | -0.851 | 0.942 |
| A29N-E28C | 431.223 | 1.106 | -5.732 | 0.962 | 1.926 | 1.034 |
| A30N-A29C | 805.425 | 0.837 | -5.327 | | 5.039 | 0.928 |
| G31N-A30C | 1251.47 | 1.007 | -4.254 | 0.894 | 8.323 | 0.822 |
| K32N-G31C | 1104.19 | 0.823 | -3.568 | 0.881 | 11.692 | 0.723 |
| T33N-K32C | - | - | -6.262 | 0.906 | 10.796 | 0.634 |
| K34N-T33C | - | - | -7.016 | 0.972 | 9.228 | 0.657 |
| E35N-K34C | 202.185 | 1.360 | - | - | 3.707 | 0.698 |
| G36N-E35C | - | - | -5.622 | - | 3.603 | 0.866 |
| V37N-G36C | - | - | -6.203 | 0.919 | 4.589 | 0.869 |
| L38N-V37C | 187.449 | 0.683 | -6.765 | 0.957 | 5.134 | 0.836 |
| T39N-L38C | 1317.285 | 0.169 | -7.943 | - | 5.703 | 0.819 |
| V40N-T39C | - | - | -5.068 | 0.921 | 6.485 | 0.801 |
| G41N-V40C | 1177.67 | 0.772 | -1.986 | 0.843 | 7.730 | 0.777 |
| S42N-G41C | - | - | -6.165 | - | 10.146 | 0.740 |
| K43N-S42C | - | - | -6.178 | - | 13.028 | 0.674 |
| T44N-K43C | 1175.190 | 0.643 | -6.111 | 0.855 | 11.849 | 0.602 |
| K45N-T44C | - | - | -7.019 | - | 10.144 | 0.630 |
| E46N-K45C | - | - | - | - | 4.628 | 0.674 |
| G47N-E46C | 513.403 | 1.735 | -5.708 | - | 4.724 | 0.835 |
| V48N-G47C | 163.944 | 1.214 | -5.986 | 0.907 | 6.269 | 0.832 |
| V49N-V48C | 140.685 | 0.692 | -7.265 | 0.928 | 8.343 | 0.783 |
| H50N-V49C | 367.473 | 1.061 | -9.486 | 0.958 | 10.519 | 0.723 |
| G51N-H50C | 885.131 | 1.966 | -6.219 | 0.916 | 7.888 | 0.664 |
| V52N-G51C | 274.331 | 0.725 | -7.068 | 0.912 | 5.462 | 0.736 |
| T53N-V52C | - | - | -6.865 | 0.954 | 4.068 | 0.808 |
| T54N-T53C | 814.16 | 0.705 | -8.041 | 0.936 | 3.021 | 0.853 |
| V55N-T54C | - | - | -8.595 | | 2.020 | 0.889 |
| A56N-V55C | 672.044 | 0726 | -7.211 | 0.924 | 0.544 | 0.924 |
| E57N-A56C | 374.917 | 0.555 | -6.535 | 1.013 | 0.075 | 0.979 |
| K58N-E57C | 629.158 | 0.691 | -6.978 | 0.984 | 4.863 | 0.997 |
| T59N-K58C | 1019.68 | 0.741 | -7.327 | 0.946 | 5.354 | 0.827 |

| | | | | | | |
|---|---|---|---|---|---|---|
| K60N-T59C | - | - | -6.965 | 0.924 | 4.215 | 0.812 |
| E61N-K60C | - | - | - | - | -1.238 | 0.849 |
| Q62N-E61N | - | - | - | - | -1.475 | 1.049 |
| V63N-Q62C | 368.242 | 0.578 | -7.443 | 1.003 | -0.774 | 1.059 |
| T64N-V63C | 1240.27 | 0.335 | -6.711 | 0.939 | -0.663 | 1.031 |
| N65N-T64C | - | - | -6.138 | - | -0.711 | 1.026 |
| V66N-N65C | 749.675 | 0.374 | -7.554 | 0.912 | -0.814 | 1.028 |
| G67N-V66C | 1047.51 | 0.867 | -6.768 | 0.826 | -0.937 | 1.032 |
| G68N-G67C | 953.059 | 1.951 | -5.397 | 0.893 | -1.066 | 1.037 |
| A69N-G68C | 961.042 | 1.037 | -5.462 | 0.912 | -1.193 | 1.042 |
| V70N-A69C | - | - | -7.941 | - | -1.312 | 1.048 |
| V71N-V70C | - | - | -8.368 | 0.929 | -1.421 | 1.052 |
| T72N-V71C | 1173.68 | 0354 | -7.319 | 0.918 | -1.513 | 1.057 |
| G73N-T72C | 1241.47 | 1.602 | -6.303 | 0.908 | -1.582 | 1.061 |
| V74N-G73C | - | - | -5.968 | 0.929 | -1.618 | 1.064 |
| T75N-V74C | 1018.64 | 0.408 | -7.354 | 0.916 | -1.602 | 1.065 |
| A76N-T75C | 1103.78 | 0.969 | -7.649 | 0.889 | -1.503 | 1.064 |
| V77N-A76C | - | - | -9.000 | 0.908 | -1.250 | 1.060 |
| A78N-V77C | 889.721 | 0.548 | -7.746 | - | -0.672 | 1.050 |
| Q79N-A78C | 197.894 | 3.911 | -8.581 | - | 0.841 | 1.026 |
| K80N-Q79C | 304.110 | 3.202 | -7.657 | 0.956 | 2.371 | 0.968 |
| T81N-K80C | 1016.72 | 0.743 | -7.686 | 0.940 | -1.276 | 0.912 |
| V82N-T81C | 296.235 | 0.719 | -7.862 | 0.991 | -5.754 | 1.051 |
| E83N-V82C | 298.986 | 0.505 | -6.862 | 1.022 | -9.394 | 1.251 |
| G84N-E83C | 1011.41 | 0.878 | -7.319 | 1.097 | -7.852 | 1.442 |
| A85N-G84C | 1078.95 | 0.924 | -5.989 | 0.949 | -6.318 | 1.358 |
| G86N-A85C | - | - | -6.349 | 0.949 | -5.707 | 1.279 |
| S87N-G86C | 1280.53 | 1.827 | -4.370 | 0.963 | -5.407 | 1.249 |
| I88N-S87C | 552.73 | 0.452 | -6.454 | 1.110 | -5.237 | 1.234 |
| A89N-I88C | 633.313 | 0.626 | -7.041 | 0.938 | -5.118 | 1.226 |
| A90N-A89C | 1009.65 | 0.667 | -7.805 | 0.936 | -5.001 | 1.221 |
| A91N-A90C | 808.213 | 0.833 | -6.905 | - | -4.839 | 1.215 |
| T92N-A91C | 1105.83 | 0.519 | -7.681 | 0.974 | -4.571 | 1.207 |
| G93N-T92C | 1361.04 | 1.462 | -5.470 | 0.927 | -4.090 | 1.195 |

| | | | | | | |
|---|---|---|---|---|---|---|
| F94N-G93C | 829.572 | 0.691 | -5.365 | - | -3.155 | 1.173 |
| V95N-F94C | 90.5266 | 1.701 | -5.324 | 0.960 | -1.001 | 1.131 |
| K96N-V95C | 483.068 | 0.921 | -6.105 | 0.996 | 1.995 | 1.040 |
| K97N-K96C | 1189.72 | 0.681 | -6.446 | - | -0.642 | 0.925 |
| D98N-K97C | 933.156 | 0.476 | -5.327 | 1.207 | -8.064 | 1.025 |
| Q99N-D98C | 633.412 | 0.806 | -6.157 | - | -9.409 | 1.369 |
| L100N-Q99C | 1143.34 | 0.245 | -6.978 | 0.974 | -9.366 | 1.443 |
| G101N-L100C | 463.91 | 1.668 | -6.068 | 1.128 | -9.104 | 1.440 |
| K102N-G101C | 468.09 | 1.941 | -6.008 | - | -9.382 | 1.426 |
| N103N-K102C | 1197.08 | 2.297 | -6.268 | 1.147 | -15.990 | 1.441 |
| E104N-N103C | 459.658 | 0.946 | -4.900 | 1.140 | -23.759 | 1.864 |
| E105N-E104C | - | - | - | - | -26.061 | 2.523 |
| G106N-E105C | - | - | -5.811 | 1.128 | -23.606 | 2.759 |
| A107N-G106C | 147.667 | 6.751 | -5.459 | 1.108 | -22.530 | 2.508 |
| P108N-A107C | 36.4415 | - | - | | -23.330 | 2.405 |
| Q109N-P108C | 202.128 | 2.201 | -7.481 | 1.121 | -25.904 | 2.481 |
| E110N-Q109C | - | - | -7.132 | 1.187 | -29.240 | 2.742 |
| G111N-E110C | 385.028 | 2.308 | -5.576 | 1.257 | -28.376 | 3.123 |
| I112N-G111C | - | - | -6.911 | 1.104 | -28.551 | 3.019 |
| L113N-I112C | 34.818 | 2.986 | -7.443 | - | -31.447 | 3.040 |
| E114N-L113C | 80.865 | 1.582 | -6.665 | - | -36.254 | 3.403 |
| D115N-E114C | 77.02 | 3.103 | -6.073 | 1.261 | -37.003 | 4.104 |
| M116N-D115C | 48.674 | 8.937 | -7.281 | - | -33.729 | 4.225 |
| P117N-M116C | 48.962 | - | - | - | -32.476 | 3.719 |
| V118N-P117C | 5.903 | 13.110 | -9.303 | 1.148 | -34.170 | 3.542 |
| D119N-V118C | 65.837 | 3.721 | -8.484 | 1.252 | -37.369 | 3.784 |
| P120N-D119C | 0.452 | - | - | - | -37.542 | 4.286 |
| D121N-P120C | 70.124 | 2.766 | -4.665 | 1.311 | -38.949 | 4.315 |
| N122N-D121C | 185.90 | 7.423 | -3.368 | 1.284 | -38.063 | 4.558 |
| E123N-N122C | 147.773 | 2.944 | -5.562 | - | -38.457 | 4.403 |
| A124N-E123C | 82.776 | 5.762 | -6.195 | 1.321 | -35.972 | 4.471 |
| T125N-A124C | 70.93 | 5.103 | -6.581 | - | -35.547 | 4.059 |
| E126N-T125C | 106.14 | 2.204 | -2.941 | 1.392 | -37.004 | 3.992 |
| M127N-E126C | - | - | -6.724 | 1.293 | -34.720 | 4.225 |

| | | | | | | |
|---|---|---|---|---|---|---|
| P128N-M127C | - | - | - | - | -33.727 | 3.866 |
| S129N-P128C | 232.832 | 3.904 | -7.565 | 1.417 | -35.611 | 3.719 |
| E130N-S129C | - | - | -6.497 | 1.371 | -39.509 | 4.002 |
| E131N-E130C | - | - | -5.765 | - | -39.422 | 4.658 |
| G132N-E131N | - | - | -6.108 | - | -35.361 | 4.642 |
| T133N-G132C | 150.157 | 3.562 | -5.484 | 1.210 | -33.352 | 3.963 |
| Q134N-T133C | 192.421 | 4.510 | -4.189 | - | -34.311 | 3.665 |
| D135N-Q134C | 154.626 | 3.466 | -4.846 | 1.541 | -36.784 | 3.805 |
| T136N-D135C | 39.900 | 5.989 | -5.995 | - | -36.235 | 4.189 |
| E137N-T136C | 56.805 | 4.119 | -5.822 | - | -36.927 | 4.101 |
| P138N-E137C | - | - | - | - | -35.355 | 4.213 |
| E139N-P138C | 56.491 | 2.124 | -6.430 | 1.266 | -35.178 | 3.962 |
| A140N-E139C | 19.876 | 0.38 | -8.170 | 1.133 | -32.659 | 3.935 |

### Figures

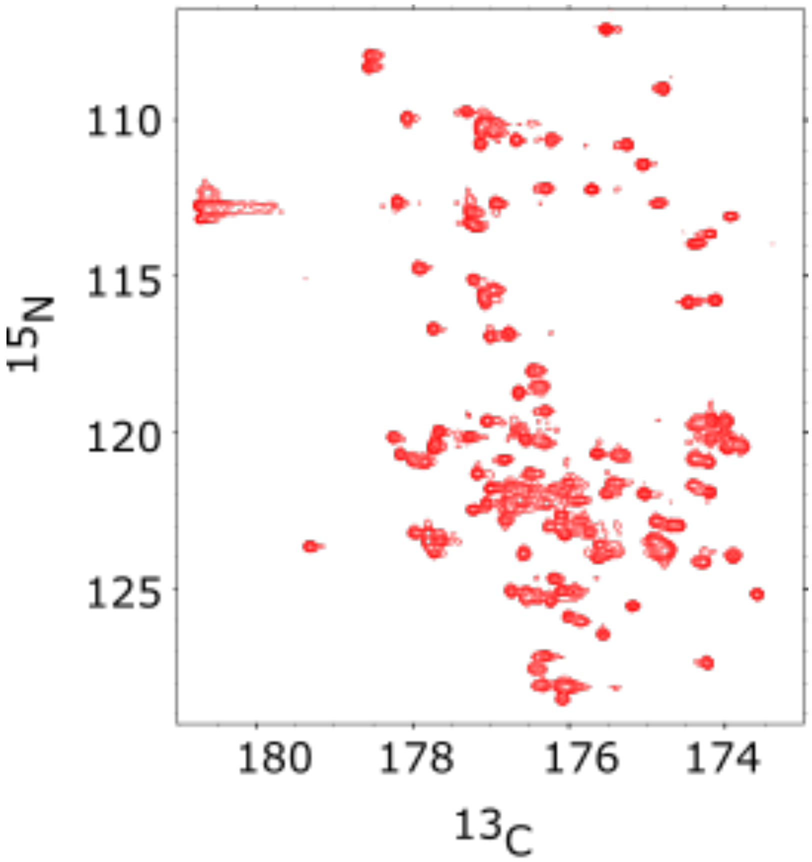


**Figure S1**. 2D CON spectrum for the measurement of hydrogen exchange for α-synuclein at 298K and pH 9.

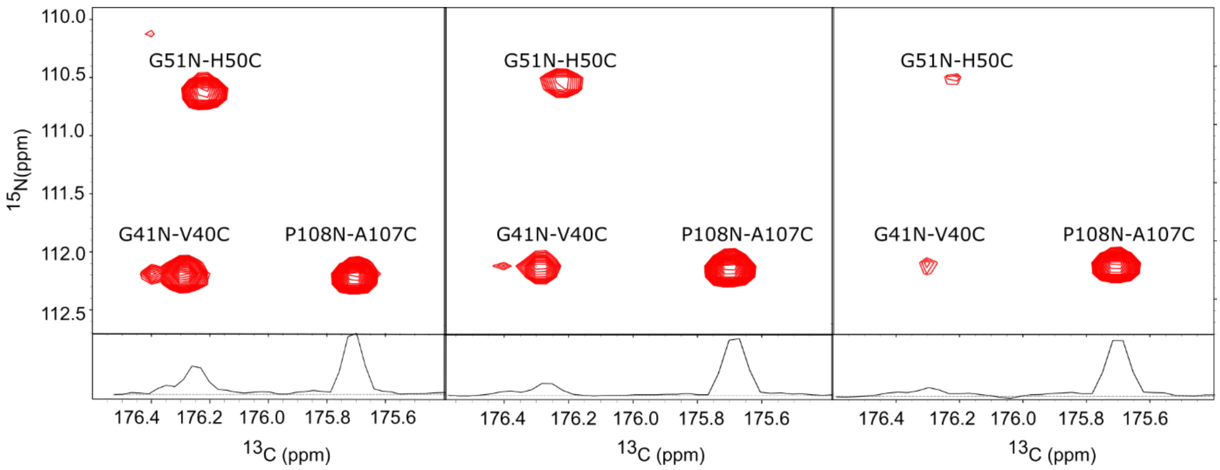


**Figure S2**. Region of the 2D CON spectrum for the measurement of hydrogen exchange obtained for reference experiment (left), with ncpmg=8 (middle) and ncpmg=2 (right). The correlation due to Pro108 is aliased in the $^{15}N$ dimension.
